## Supplemental Data for "A concerted increase in readthrough and intron retention drives transposon expression during aging and senescence"

| **Dataset** | **Transp. family** | **up** | **down** | **Fraction up** | **mean log2 fc** |
| --- | --- | --- | --- | --- | --- |
| Aging | LTR | 770 | 163 | 0.825 | 0.328 |
|  | DNA | 845 | 70 | 0.923 | 0.430 |
|  | LINE | 3337 | 352 | 0.905 | 0.408 |
|  | SINE | 1518 | 589 | 0.720 | 0.326 |
|  | **Transp. family** | **up** | **down** | **Fraction up** | **mean log2 fc** |
| Senescence | LTR | 1640 | 183 | 0.900 | 2.445 |
|  | DNA | 611 | 149 | 0.804 | 1.766 |
|  | LINE | 2901 | 481 | 0.858 | 2.097 |
|  | SINE | 1960 | 596 | 0.767 | 1.581 |

**Table S1**
Numbers of significantly up- and downregulated transposon families with aging and cellular senescence. Mean log2-fold change (“log2 fc”) and fraction upregulated is also shown. Aging data from Fleischer et al. (2018) and senescence data from Colombo et al. (2018).

| **Aging** | sig. readthrough | no sig. readthrough | Fraction |  |
| --- | --- | --- | --- | --- |
| transposon | 579 | 63203 | 0.009 |  |
| sig. transposon | 158 | 7535 | 0.021 | p<0.0001 |
| **Senescence** |  |  |  |  |
| transposon | 1840 | 122333 | 0.015 |  |
| sig. transposon | 574 | 8156 | 0.070 | p<0.0001 |
| **Aging** | any. readthrough | no readthrough | Fraction |  |
| transposon | 1357 | 62425 | 0.022 |  |
| sig. transposon | 265 | 7428 | 0.036 | p<0.0001 |
| **Senescence** |  |  |  |  |
| transposon | 6061 | 118112 | 0.051 |  |
| sig. transposon | 751 | 7979 | 0.094 | p<0.0001 |

**Table S2**
Number of transposons and significant transposons that are located in readthrough regions. p-value by Fisher’s exact test.

| **Dataset** | **Analysis** | **Transposons** | **R** | **n** | **P-value** |
| --- | --- | --- | --- | --- | --- |
| Aging  Fleischer et al. | sig vs sig | downstream | 0.33 | 156 | p<0.0001 |
|  | sig vs sig | any | 0.11 | 465 | p=0.022 |
|  | all vs all | downstream | 0.29 | 1337 | p<0.0001 |
|  | all vs all | all | 0.13 | 9418 | p<0.0001 |
| Senescence  Colombo et al. | sig vs sig | downstream | 0.88 | 562 | p<0.0001 |
|  | sig vs sig | any | 0.74 | 1246 | p<0.0001 |
|  | all vs all | downstream | 0.76 | 5971 | p<0.0001 |
|  | all vs all | all | 0.45 | 33232 | p<0.0001 |

**Table S3**
Correlation (R) between the expression levels of transposons and readthrough at the adjacent gene. We included either all significant elements in this analysis (“sig vs sig”) or all expressed elements (“all vs all”). Transposons in the “any” category can be located upstream, intragenic, intronic or downstream to the gene and its readthrough region.

| **Model** | **Adjusted R-squared** |
| --- | --- |
| transposon ~ intron | 0.405 |
| transposon ~ intron + readthrough | 0.562 |

**Table S4**
Comparison of two linear models to predict normalized transposon expression in the aging dataset using either normalized intron retention levels or also including normalized readthrough levels. P<0.0001 by ANOVA.

| **Senescence** | LINE1 | non LINE1 | | **Aging** | LINE1 | non LINE1 |
| --- | --- | --- | --- | --- | --- | --- |
| intragenic | 63.9 | 59.0 |  | intragenic | 80.4 | 87.6 |
| intronic | 53.4 | 46.2 |  | intronic | 52.1 | 38.2 |
| downstream | 13.1 | 20.4 |  | downstream | 15.3 | 9.0 |
| upstream | 12.2 | 12.0 |  | upstream | 3.6 | 3.1 |
| sum | 89.2 | 91.4 |  | sum | 99.3 | 99.6 |

**Table S5**
Percentage of transposons that are intragenic, intronic, downstream or upstream of genes (within 25kb) and the sum total of intragenic, downstream and upstream.

| **Transposon class** | **Aging** | **Senescence** |
| --- | --- | --- |
| active, significant LINE-1 | 0.65 | 0.97 |
| significant LINE-1 | 0.94 | 0.88 |
| significant transposon | 0.84 | 0.84 |
| any transposon | 0.61 | 0.62 |

**Table S6**
The fraction of transposons upregulated with aging or cellular senescence in each class.

| **Dataset** | **Group** | **Transposons** | **any L1** | **L1 w/**  **ORF1p** | **P-value** | **L1 w/ ORF2p** | **P-value** |
| --- | --- | --- | --- | --- | --- | --- | --- |
| Aging  Fleischer et al. | outliers | 283 | 91 | 0 | NS | 0 | NA |
|  | sig | 7693 | 2820 | 8 | <0.05 | 0 | NA |
|  | all | 63782 | 12214 | 30 | NA | 2 | NA |
| Senescence  Colombo et al. | outliers | 265 | 86 | 2 | <0.05 | 0 | NA |
|  | sig | 8730 | 2506 | 19 | <0.0001 | 2 | NS |
|  | all | 124173 | 27167 | 95 | NA | 24 | NA |

**Table S7**
Numbers of ORF1p- and ORF2p-encoding LINE-1 elements among all expressed transposons, transposons significantly changed with aging and transposons with higher-than-expected expression during aging (“outliers”). The p-value was determined by comparison with randomly sampled elements from the “all” group.

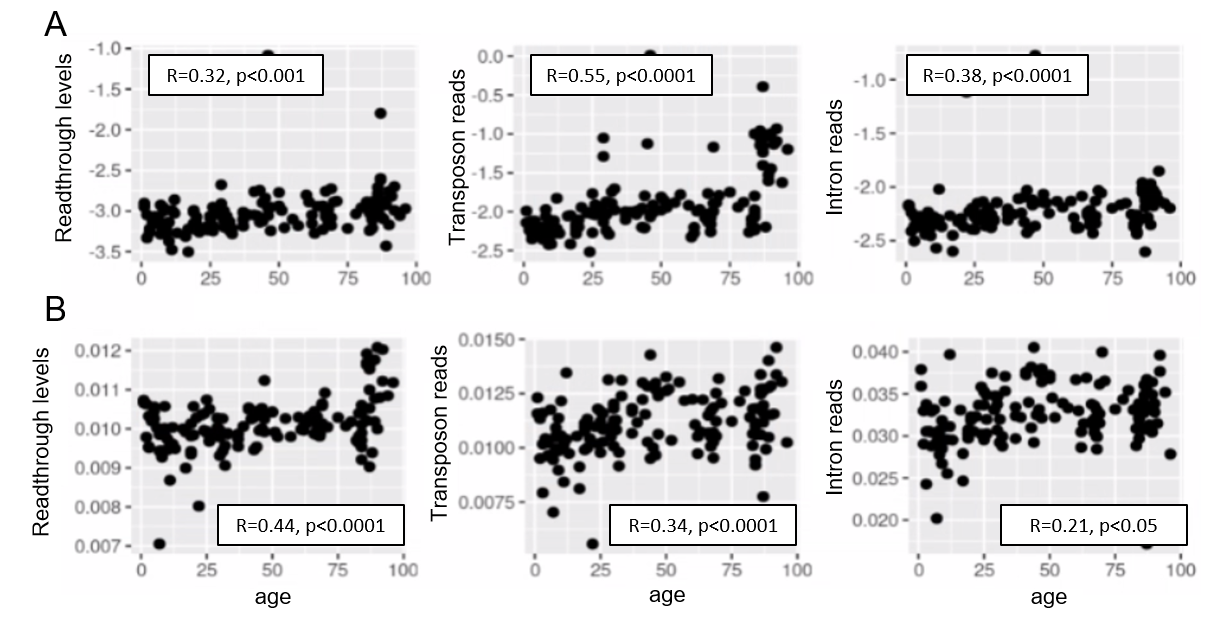

**Fig. S1. Aging promotes readthrough, transposon expression and intron retention.**
Readthrough, transposon expression and intron retention are elevated with aging when the expression of each element is normalized to the nearest gene (A) and similarly when all the reads are normalized to library size (B). Reads from the top 1000 differentially expressed elements were used in this analysis (n<1000 for readthrough after filtering).

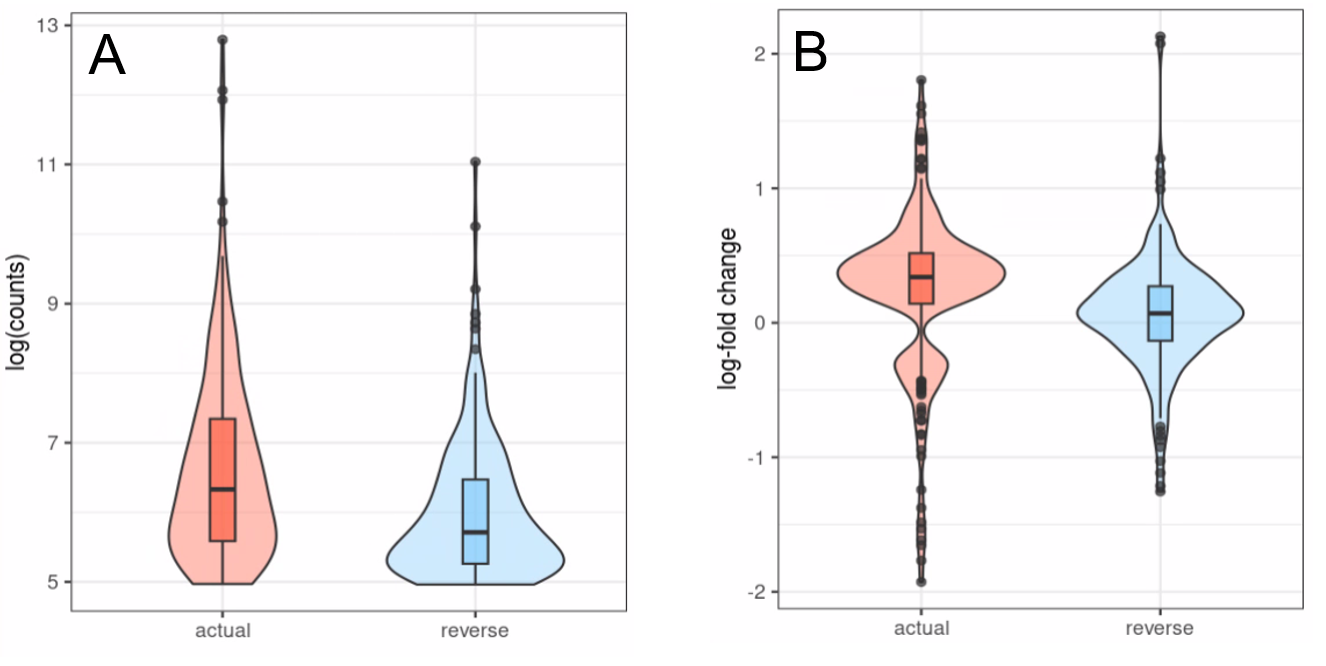

**Fig. S2. Transposons show strand-specific expression.**
Transposons with inverted strandedness (“reverse”) show lower expression levels (log counts; A) and no differential expression with age (B) when compared to matched differentially expressed transposons (“actual”). For this analysis we selected all transposons showing significant differential expression with age in the actual dataset that also showed at least minimal expression in the strand-inverted analysis (n=226). Data from Fleischer et al. (2018).
(A) The log (counts) are clipped because we only used transposons that passed minimal read filtering in this analysis.
(B) The distribution of expression values in the actual dataset is bimodal and positive since some transposons are significantly up- or downregulated. This bimodal distribution is lost in the strand-inverted analysis.

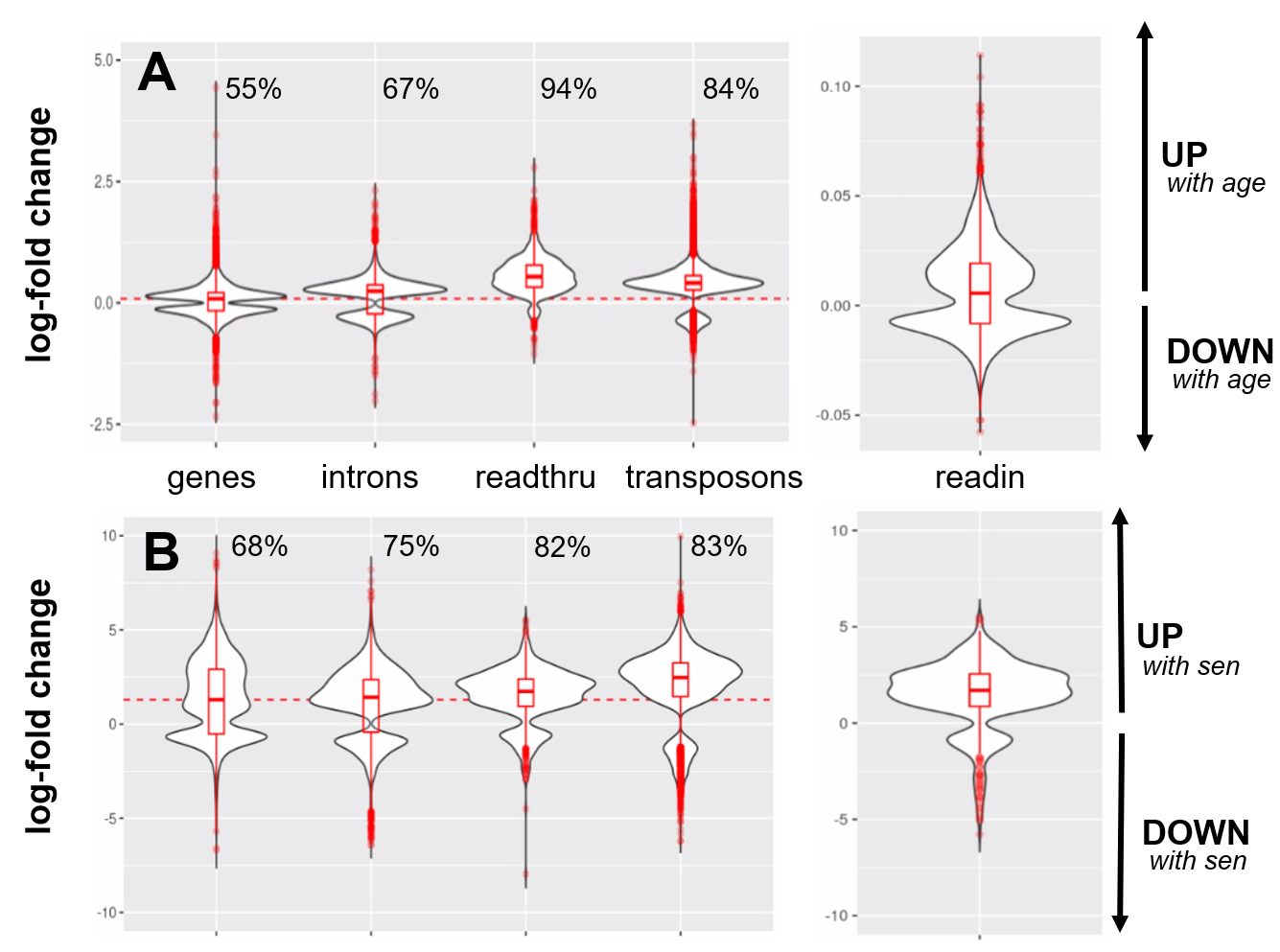
 **Fig. S3. Stronger upregulation of transposons and introns than of genic transcripts.**
Intron, readthrough and transposon elements are more strongly elevated with age (A) and cellular senescence (B) than are genic transcripts. The percentage of upregulated transcripts is indicated above each violin plot and the median log10-fold change for genic transcripts is indicated with a dashed red line.
(A) Log10-fold changes plotted for all elements that significantly change with age (genes, introns, readthrough, transposon and read-in transcripts).
(B) Log10-fold changes plotted for all elements that significantly change with induced senescence (genes, introns, readthrough, transposon and read-in transcripts). The four senescent conditions (H2O2, 5-Aza, Adriamycin, replicative senescence) are grouped together and compared with four control conditions (serum-starved, immortalized, intermediate passage and early passage)

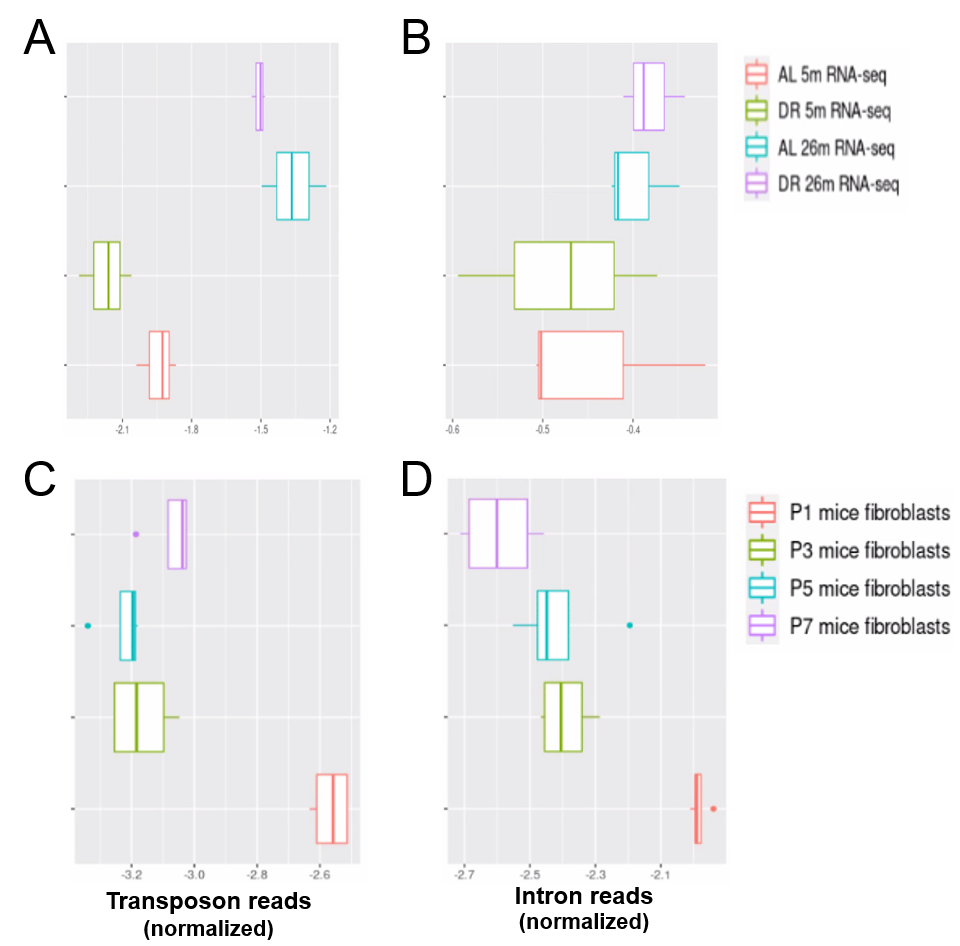

**Fig. S4. Increased transposon expression and intron retention in liver of aged mice.**
Transposon expression (A) and intron retention (B) are increased in the liver of 26 month-old mice. In contrast, transposon expression (C) and intron retention (D) are decreased with replicative senescence of mouse fibroblasts. Normalized counts from all (A, B) and top 1000 (C, D) differentially expressed genes, transposons and introns were used in this analysis. Mouse liver data is from Hahn et al. 2017 (GSE92486) and mouse replicative senescence data from Wang et al. 2022 (GSE179880).
A) Transposon reads normalized by the expression of adjacent genes plotted for each sample (ad libitum [AL] 5 month-old, AL 26 month-old, dietary restricted [DR] 5 month-old and DR 26 month-old, n=3 per group)
B) Intron reads normalized by the expression of adjacent genes plotted for each sample as in (A).
C) Transposon reads normalized by the expression of adjacent genes plotted for each sample (fibroblasts from passage [P] 1, 3, 5 and 7, n=4 per group)
D) Intron reads normalized by the expression of adjacent genes plotted for each sample as in (C).

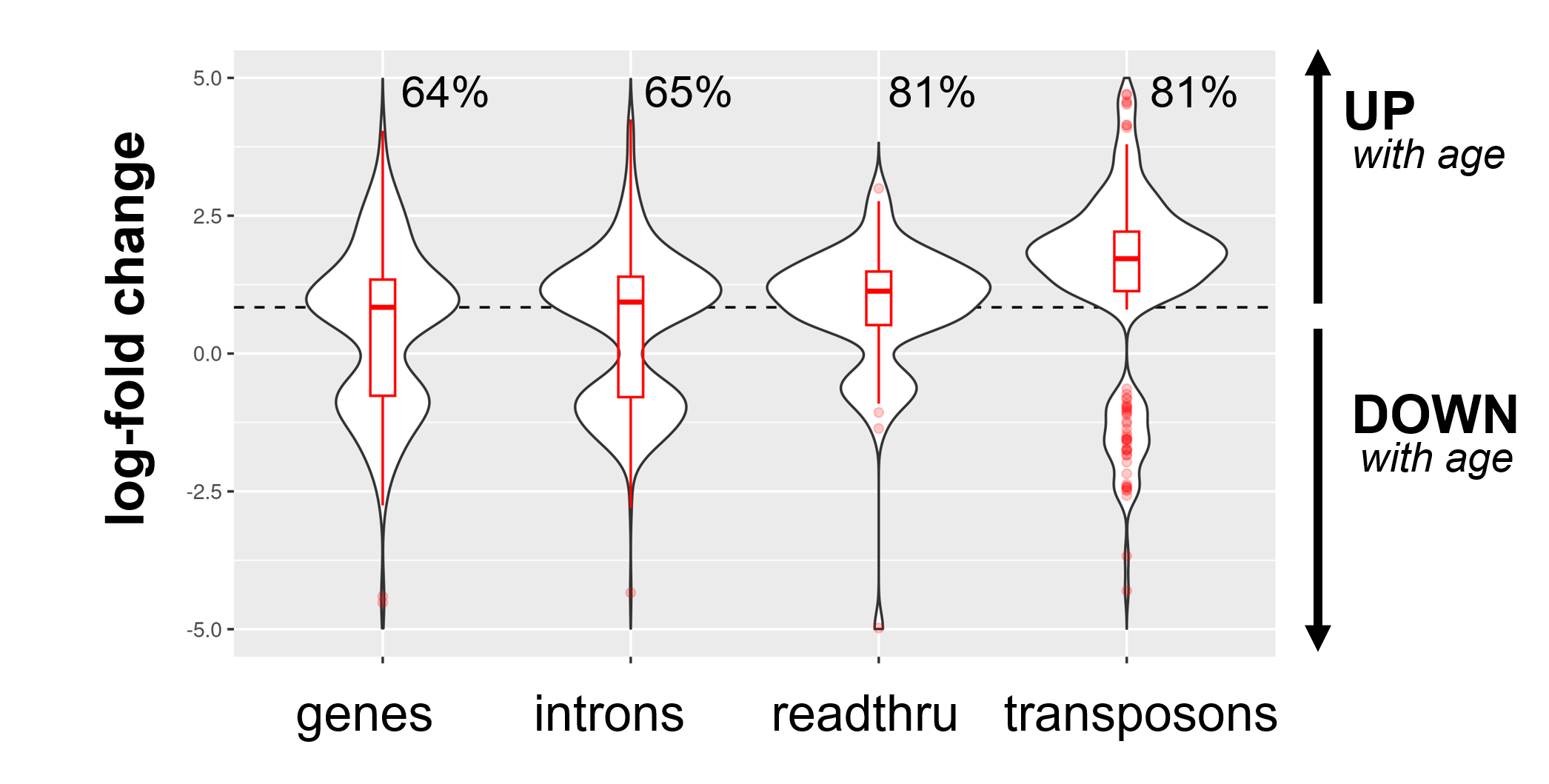

**Fig. S5. Aging promotes readthrough, transposon expression and intron retention in mouse liver.**
Intron, readthrough and transposon elements are elevated in the liver of aging mice (26 vs 5-month-old, n=6 per group). Readthrough and transposon expression is especially elevated even when compered to genic transcripts. The percentage of upregulated transcripts is indicated above each violin plot and the median log10-fold change for genic transcripts is indicated with a dashed red line.

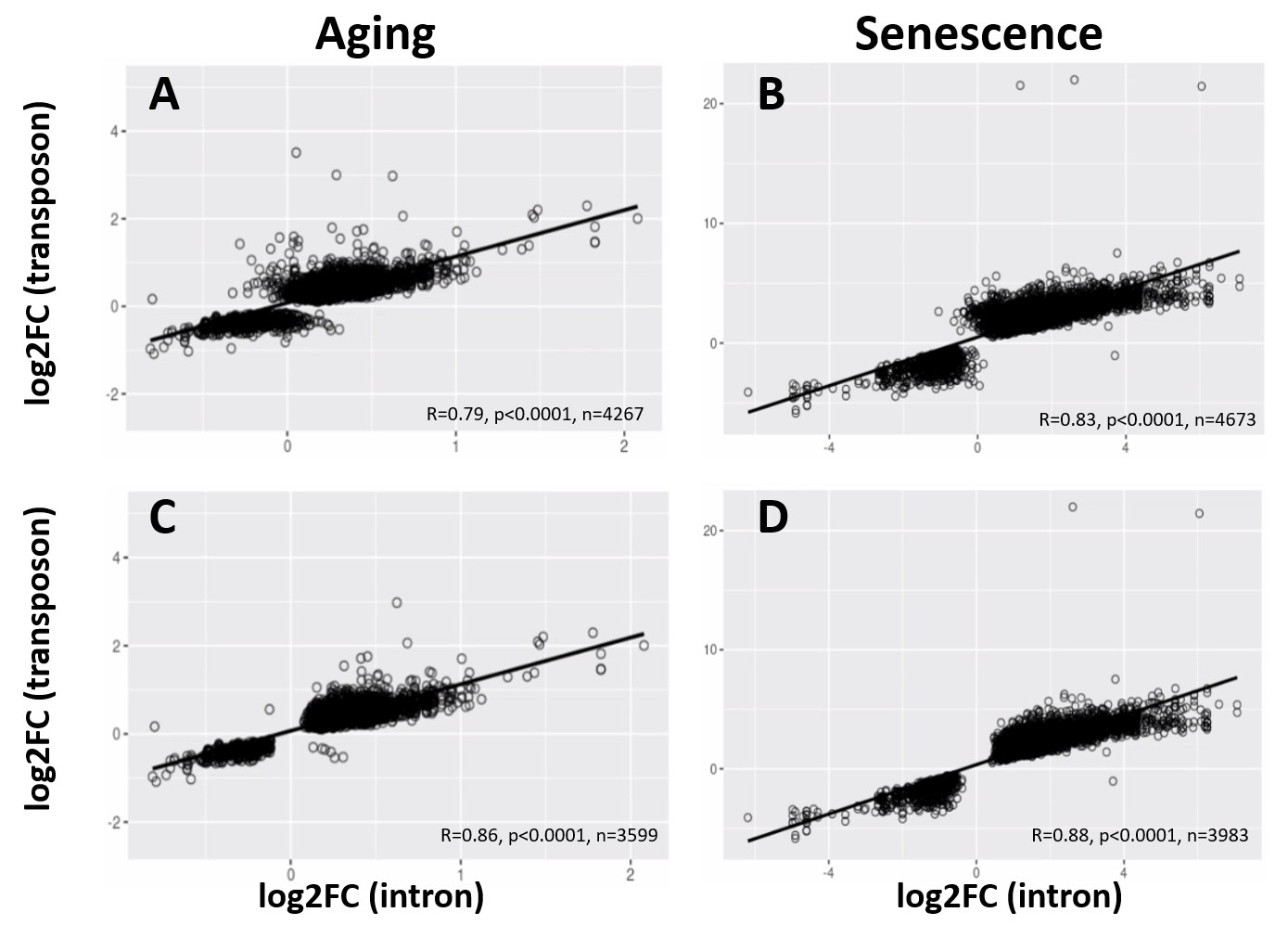

**Fig. S6. Correlated age-related increase of transposon and intron loci.**
Intron and transposon expression show a significant correlation. For every transposon within an intron, the correlation between the log-fold change of the intron and the transposon is plotted. Scatter plots are shown for all transposons and introns (A, B) and all transposons and introns that are significantly changed (C, D) in the aging dataset (A, C) and with cellular senescence (B, D).

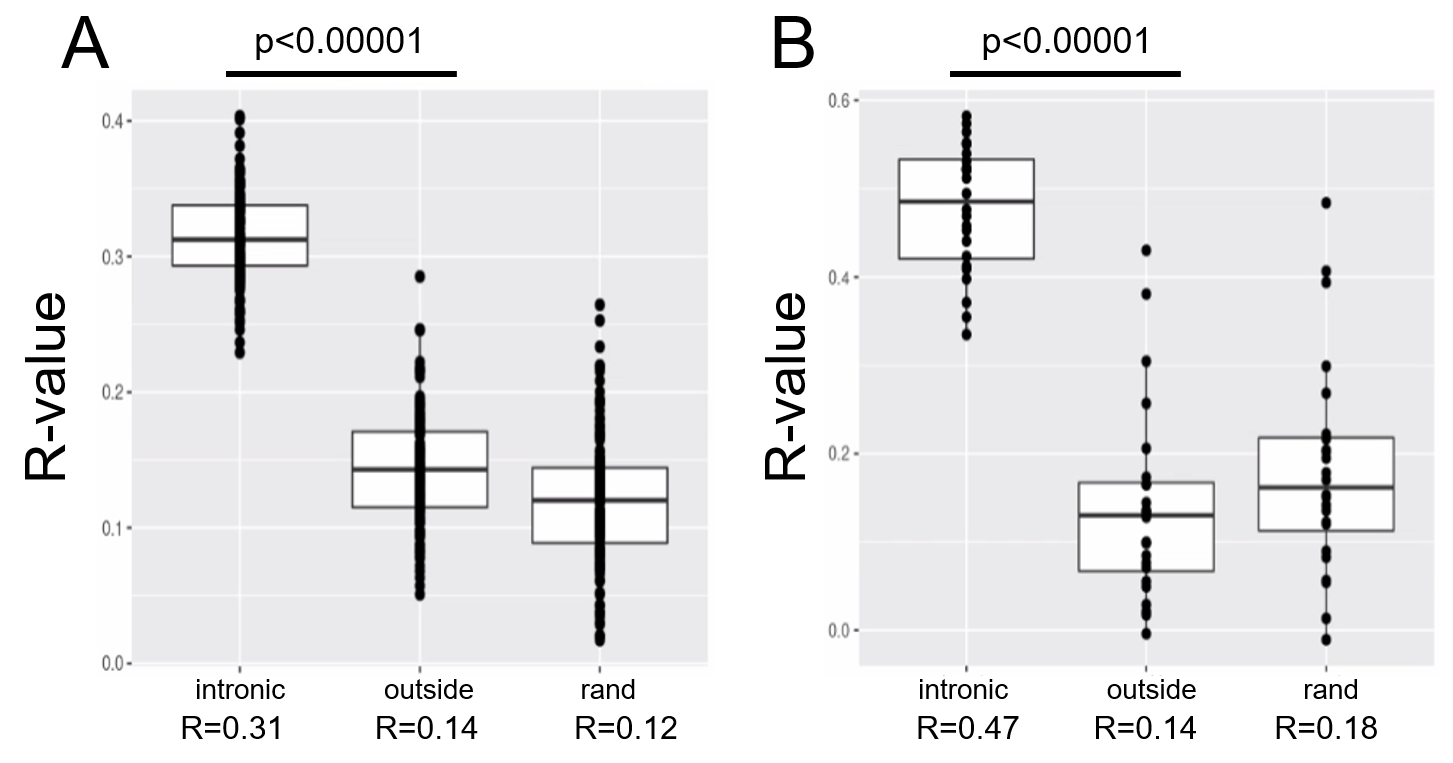

**Fig. S7. Correlated expression of transposon and intron loci.**
Intronic counts are correlated with the counts of intronic transposons in the aging dataset (A) and in the senescence dataset (B). In contrast, intragenic reads aligning outside of introns do not correlate with the counts of intronic transposons (labeled “outside”). Intronic counts also show no correlation with randomized transposon counts (labeled “rand”). Pearson correlation was performed for each sample in these datasets and the R-values for all the samples are shown here. Intronic counts for a gene were defined as the sum of all its intronic counts.

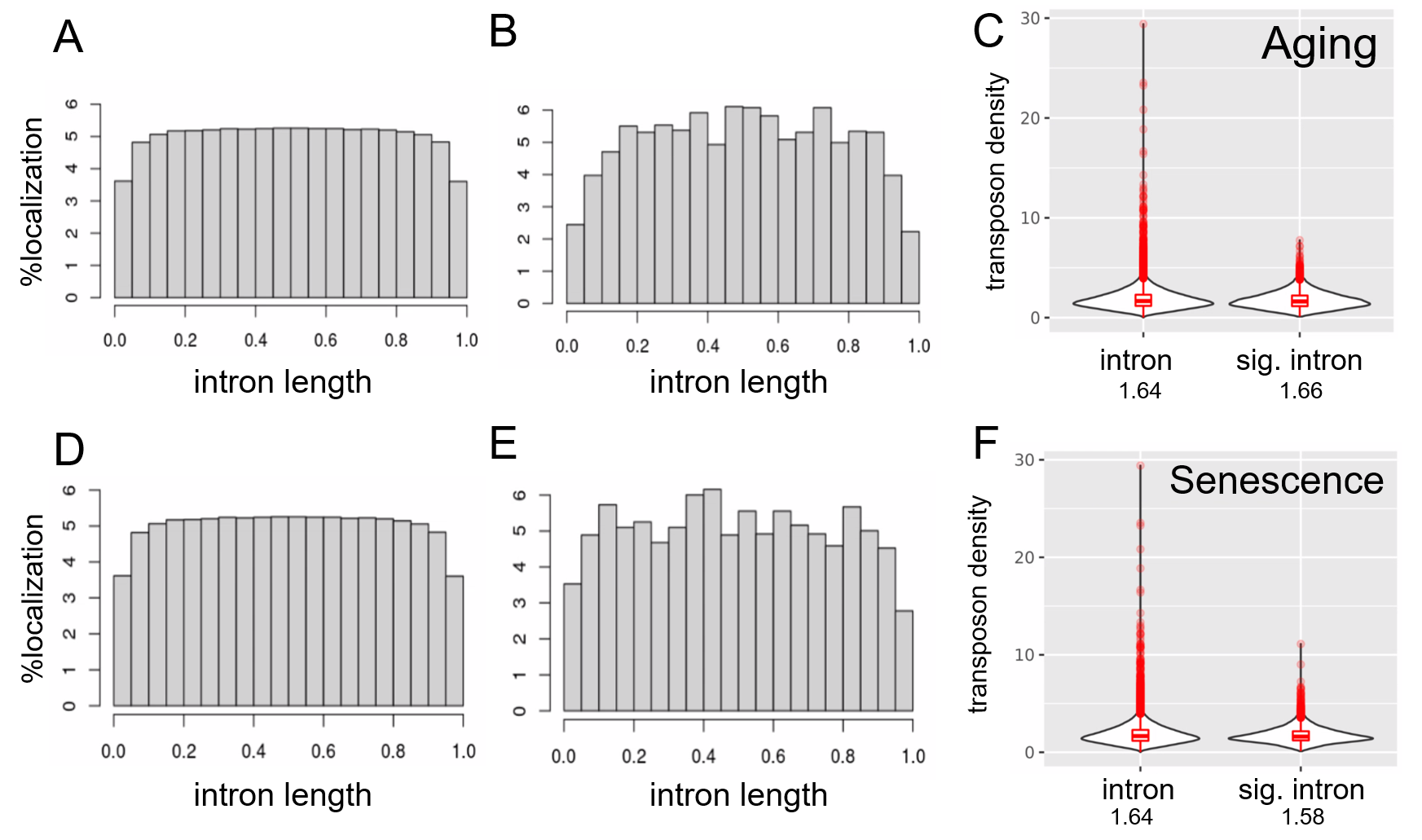
**Fig. S8. Transposons are depleted at splice junctions.**
Transposons are evenly distributed within introns except for the region close to splice junctions (A-E). Transposons appear to be excluded from the splice junction-adjacent region both in all introns (A, D) and in significantly retained introns (B, E). In addition, transposon density of all introns and significantly retained introns is comparable (C, F). We included only introns containing at least one transposon in this analysis and normalized their length to 1.
A) Distribution of 2292769 transposons within 163498 introns among all annotated transposons.
B) Distribution of 195190 transposons within 14100 introns significantly retained with age.
C) Density (transposon/1kb of intron) of transposons in all introns (n=163498) compared to significantly retained introns (n=14100).
D) as in (A)
E) Distribution of 428130 transposons within 13205 introns significantly retained with induced senescence.
F) Density (transposon/1kb of intron) of transposons in all introns (n=163498) compared to significantly retained introns (n=13205).

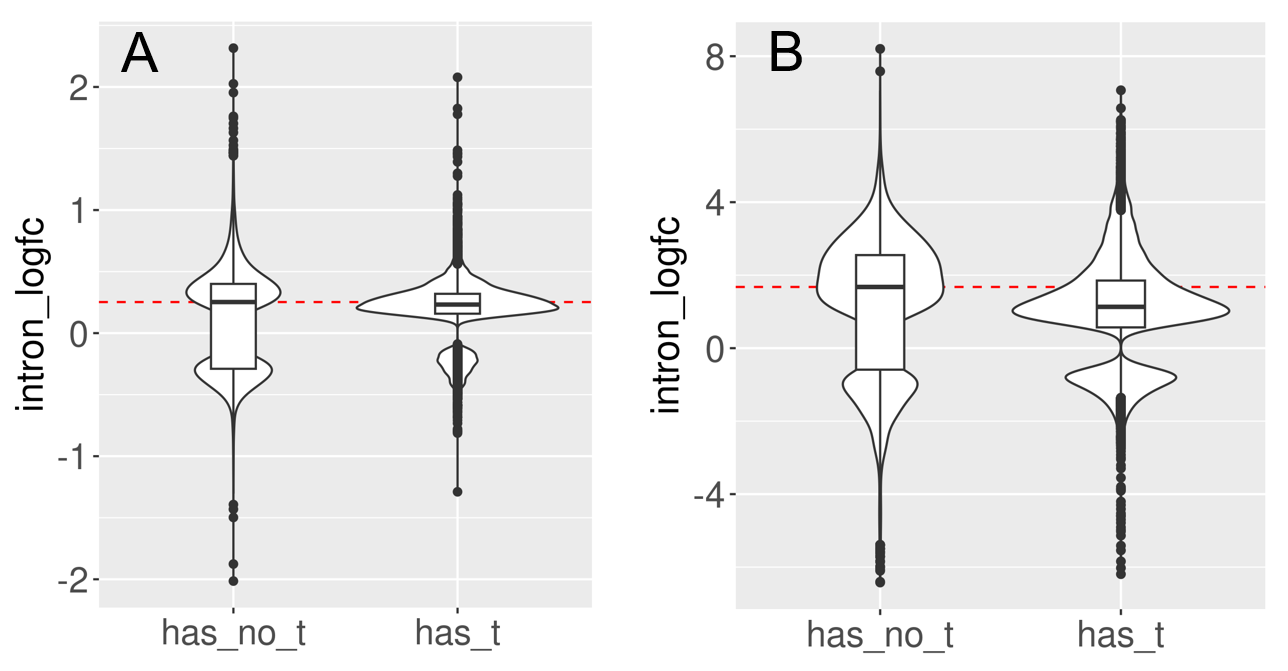

**Fig. S9. Introns with and without transposons are retained during aging and senescence.**
We split the set of introns that significantly change with cellular aging (A) or cell senescence (B) into introns that contain at least one transposon (has_t) and those that do not contain any transposons (has_no_t). Intron retention is increased in both groups. In this analysis we included all introns that passed minimal read filtering (n=63782 in A and n=124173 in B). Median log-fold change indicated with a dashed red line for the group of introns without transposons.

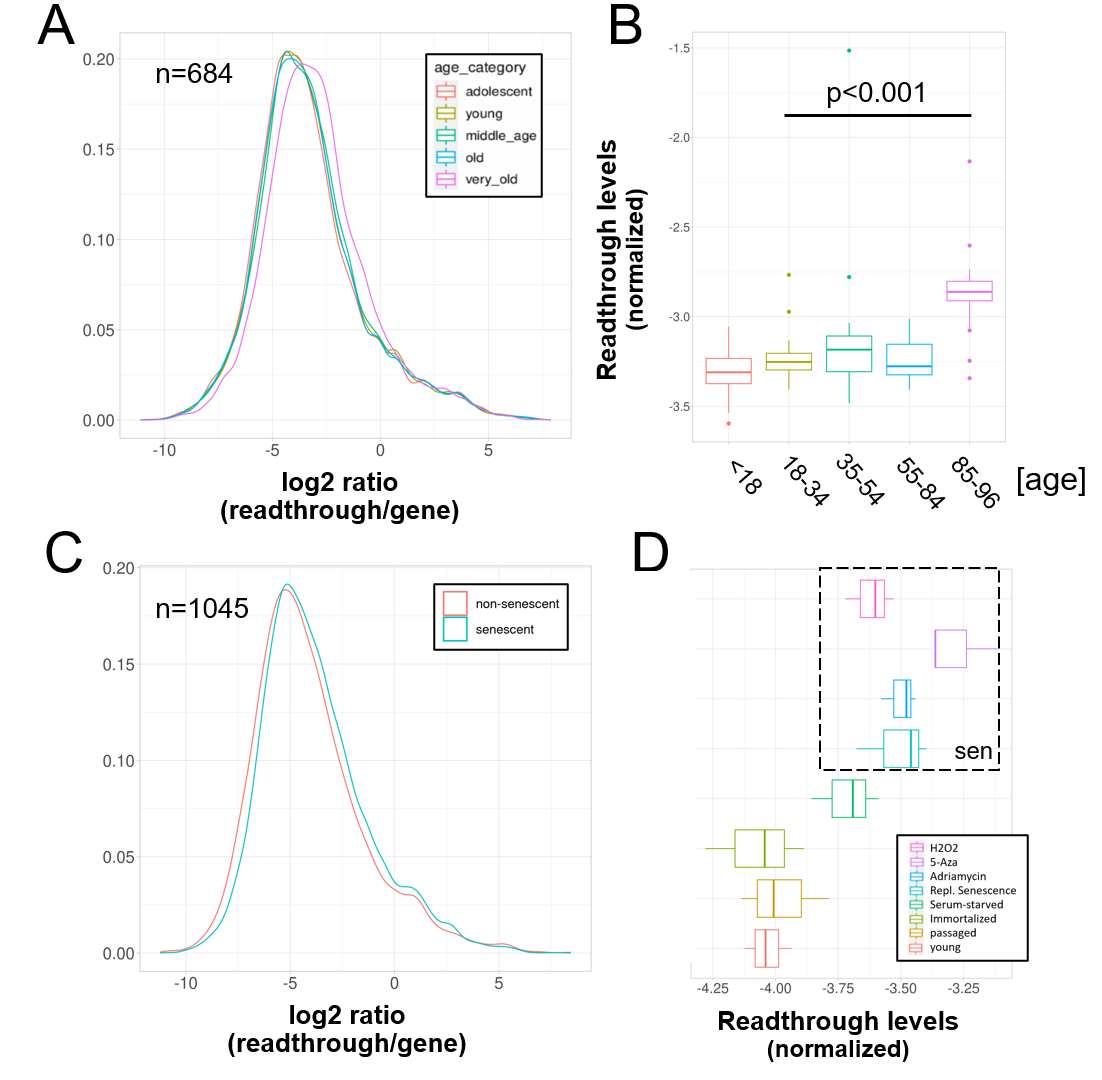

**Fig. S10. Increased readthrough levels with aging and senescence.**
Readthrough transcription is increased in fibroblasts isolated from the very old (A, B) and after induction of senescence in vitro (C, D).
A) Readthrough was determined in a region 0 to 10 kb downstream of genes for a subset of genes that were at least 10 kb away from the nearest neighboring gene (n=684 regions). The log2 ratio of readthrough to gene expression is plotted across five age groups (adolescent n=32, young n=31, middle-aged n=22, old n=37 and very old n=21).
B) As in (A) but data is plotted on a per sample basis.
C) Readthrough was determined in a region 0 to 10 kb downstream of genes for a subset of genes that were at least 10 kb away from the nearest neighboring gene (n=1045 regions). The log2 ratio of readthrough to gene expression is plotted for the groups comprising senescence (n=12) and the non-senescent group (n=6).
D) As in (D) but data is plotted on a per sample basis and for additional control datasets (serum-starved, immortalized, intermediate passage and early passage). N=3 per group.

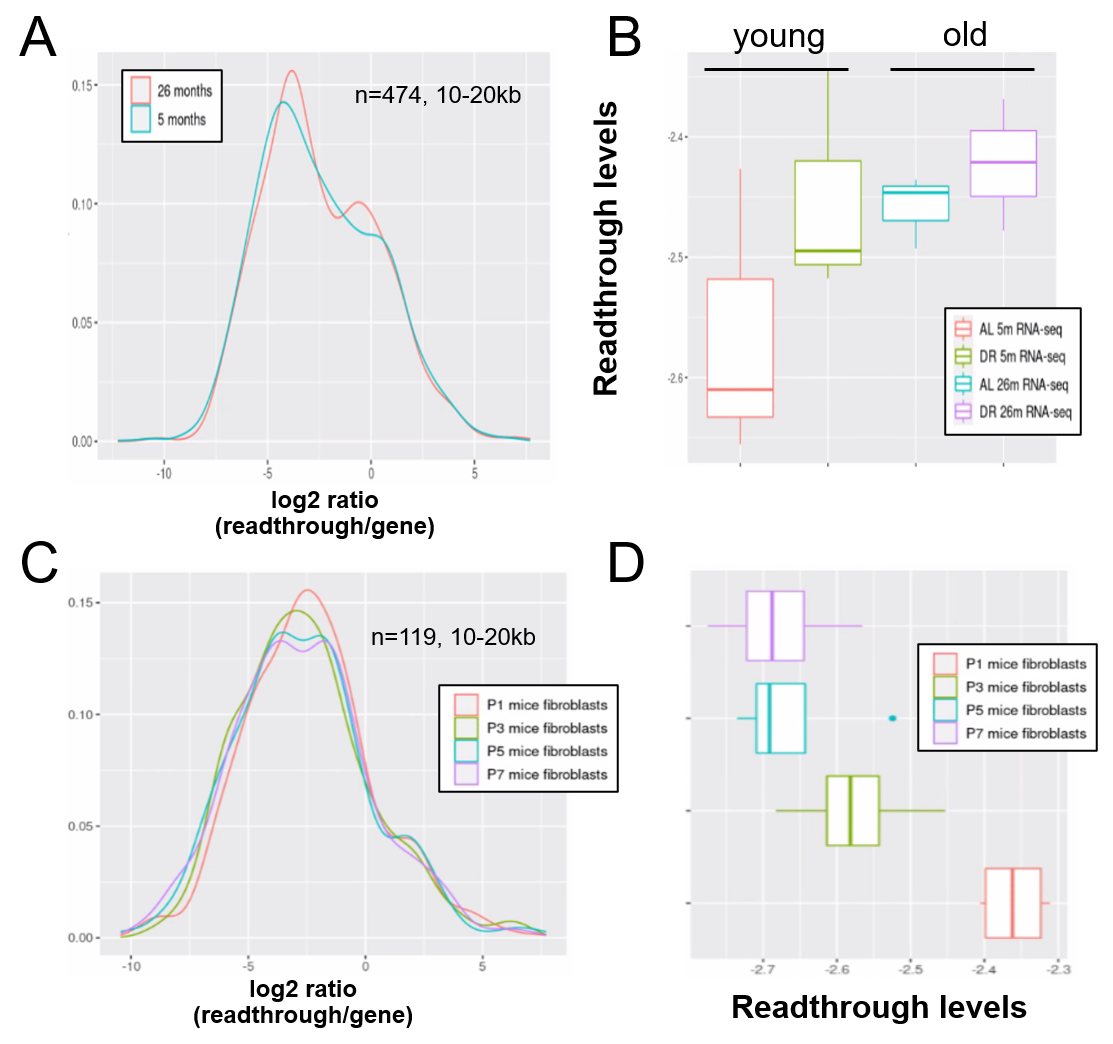
 **Fig. S11. Readthrough transcription might be increased with age in mouse liver, but not during in vitro senescence of mouse fibroblasts.**
Readthrough transcription is non-significantly increased with age in mouse liver (A, B) and this increase is not attenuated by dietary restriction (B). In contrast, there is no increase in readthrough when mouse fibroblasts undergo replicative senescence (C, D).
A) Readthrough was determined in a region 10 to 20kb downstream of genes for a subset of genes that were at least 20kb away from the nearest neighboring gene (n=474 genes). The log2 ratio of readthrough to gene expression is plotted for liver samples from 5- and 26-month-old mice (n=6 per group).
B) As in (A) but data is plotted on a per-sample-basis (n=3 per group).
C) Readthrough was determined in a region 10 to 20kb downstream of genes for a subset of genes that were at least 20kb away from the nearest neighboring gene (n=119 genes). The log2 ratio of readthrough to gene expression is plotted across five age groups (n=4 per group).
D) As in (C) but data is plotted on a per-sample-basis.

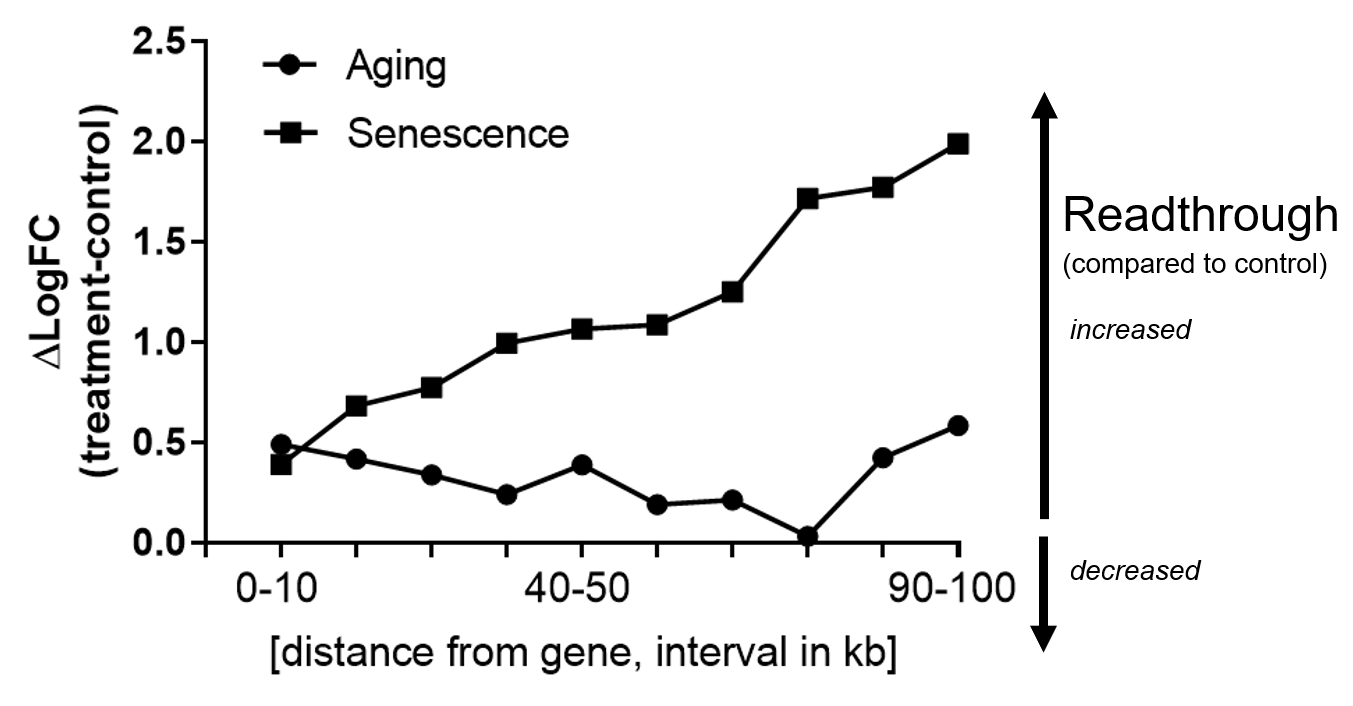

**Fig. S12. Readthrough is increased downstream of genes over a wide range of distances.**
Transcriptional readthrough observed with aging or cellular senescence is elevated over a wide region downstream of genes. Readthrough levels were determined for each 10kb interval downstream of genes, in the range between 0 and 100kb using the same approach as in Fig. 4. Values for each gene and treatment group were log-transformed and pooled. The difference between treatment (either senescence or aging) and controls is shown here. Higher numbers indicate stronger readthrough. The senescence group comprises data from H2O2 treatment, Adriamycin, 5-Azacytidine and replicative senescence. The aging group comprises data from the very old subgroup (85 to 96 years-old) and was compared to the middle-aged group.

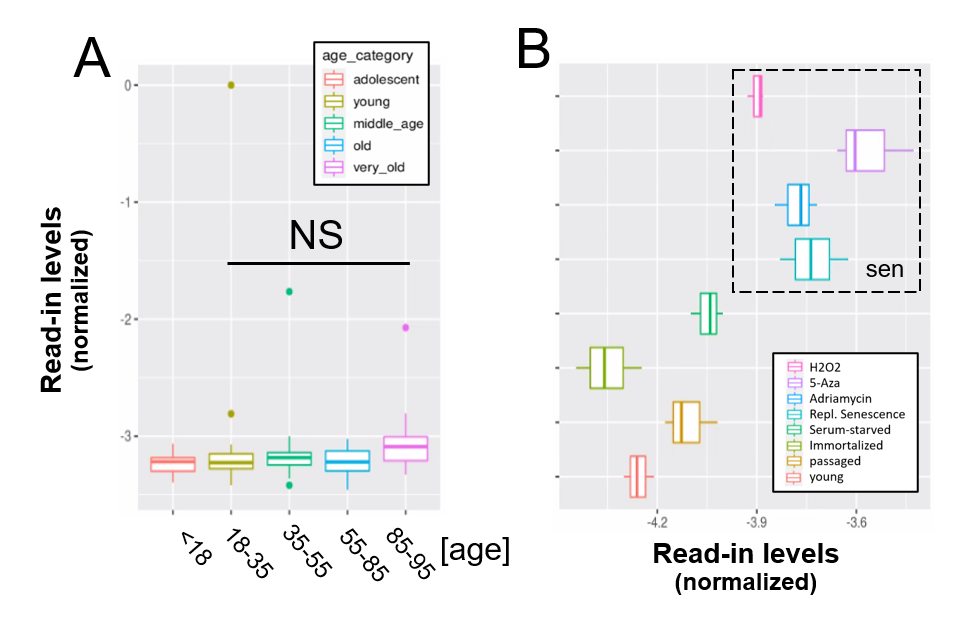
 **Fig. S13. Read-in does not change with age and is increased by senescence.**
Read-in expression is unchanged with aging (A) whereas it is increased with cellular senescence (B).
A) Read-in normalized by the expression of adjacent genes plotted across five age groups (adolescent n=32, young n=31, middle-aged n=22, old n=37 and very old n=21).
B) Read-in normalized by the expression of adjacent genes plotted across four senescent conditions (H2O2, 5-azacytidine, adriamycin, replicative senescence) and four other conditions (serum-starved, immortalized, intermediate passage and early passage). N=3 per group.

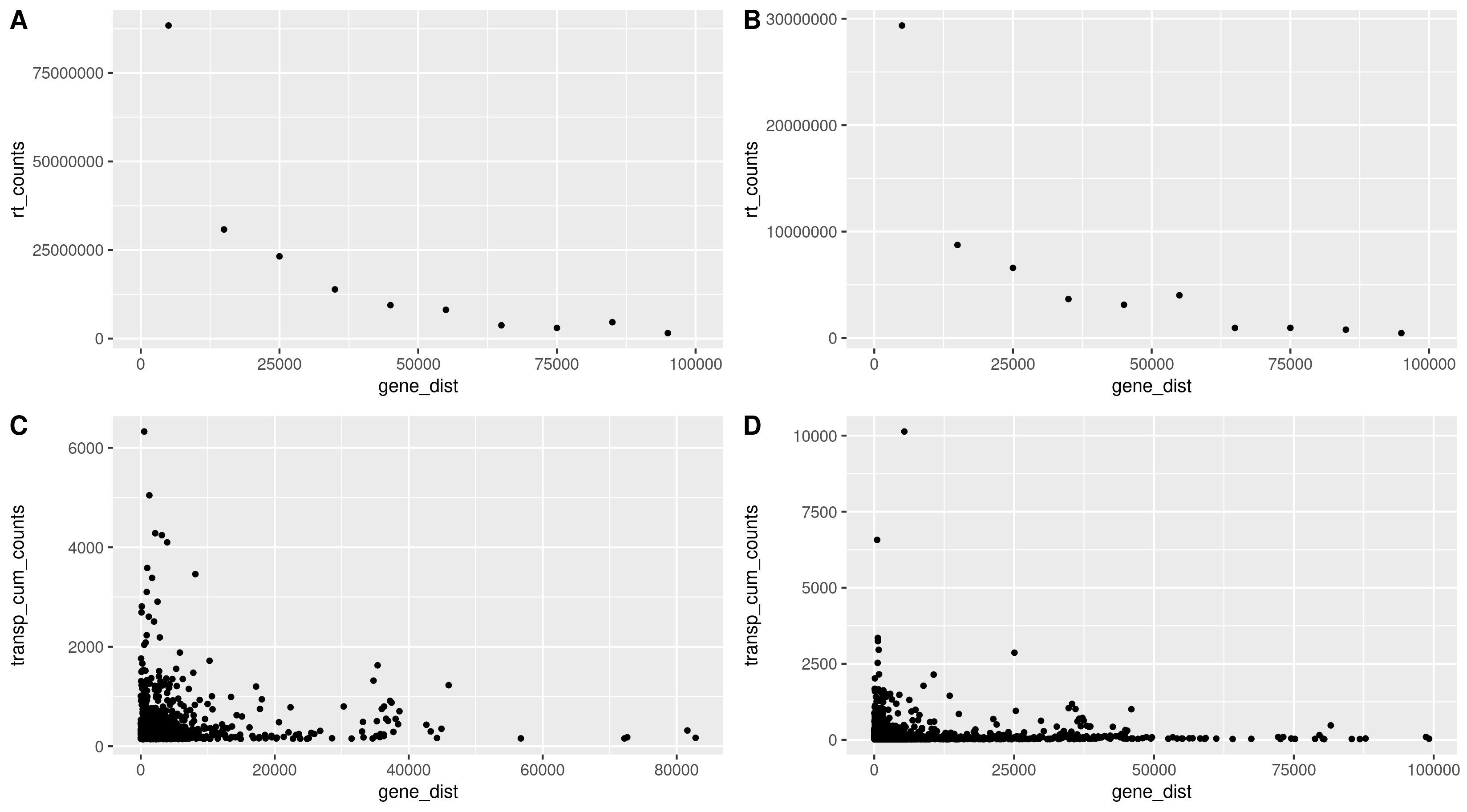

**Fig. S14. Readthrough and transposon counts are higher closer to genes.**
Readthrough counts (rt_counts) decrease exponentially downstream of genes, both in the aging dataset (A) and in the cellular senescence dataset (B). Although noisier, the pattern for transposon counts (transp_cum_counts) is similar with higher counts closer to gene terminals, both in the aging dataset (C) and in the cellular senescence dataset (D). Readthrough counts are the cumulative counts across all genes and samples. Readthrough was determined in 10 kb bins and the values are assigned to the midpoint of the bin for easier plotting. Transposon counts are the cumulative counts across all samples for each transposon that did not overlap a neighboring gene. n=801 in (C) and n=3479 in (D).

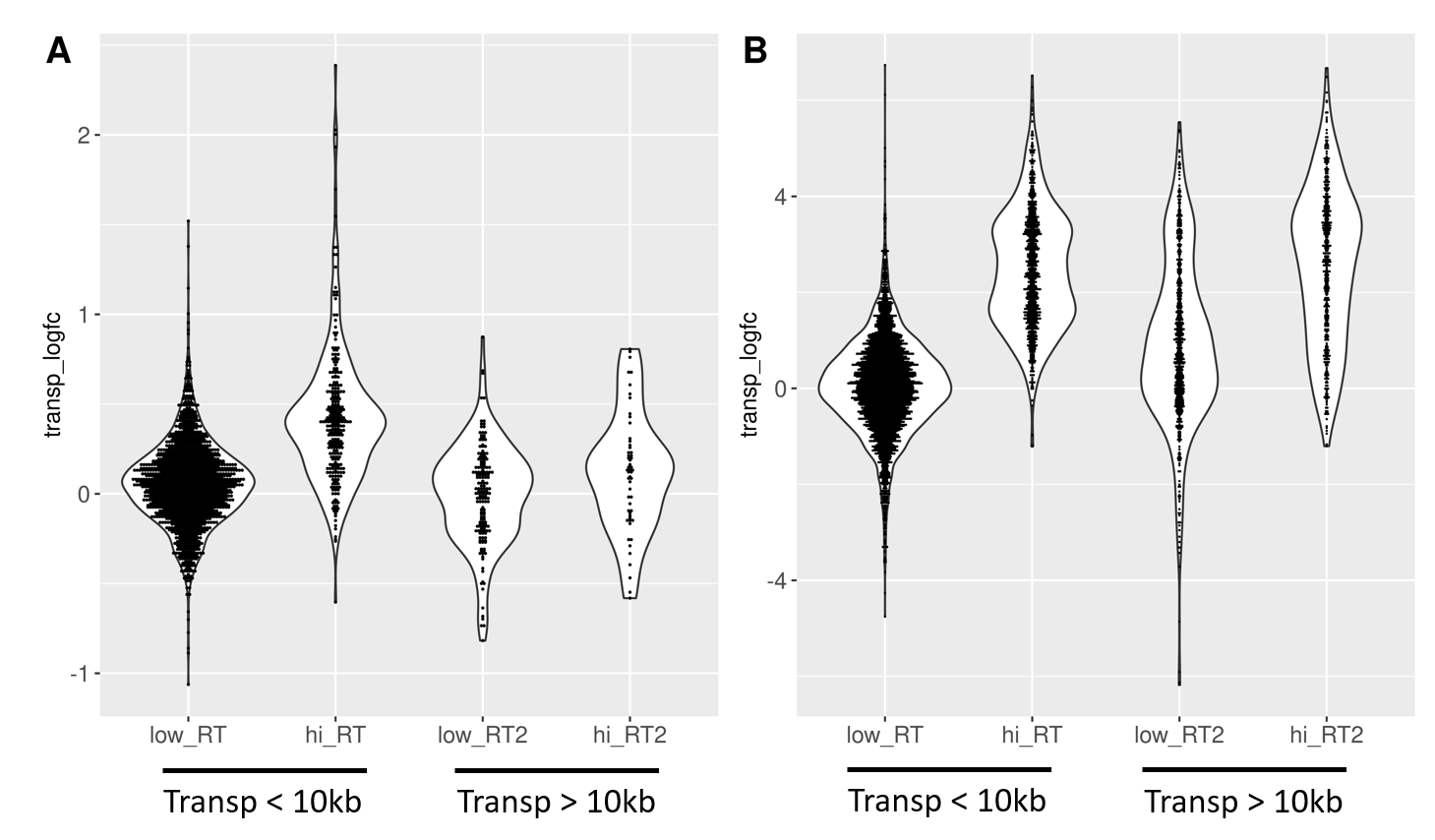

**Fig. S15. Age-related transposon expression downstream of high readthrough genes is elevated.**
Transposons found downstream of genes with high readthrough (hi_RT) show a more pronounced log-fold change (transp_logfc) than transposons downstream of genes with low readthrough (low_RT). This is true in fibroblasts isolated from aged donors (A) and with cellular senescence (B). Furthermore, the difference between high and low readthrough region transposons is diminished for transposons that are more than 10 kb downstream of genes (“Transp > 10 kb”). Transposons in high readthrough regions were defined as those in the top 20% of readthrough log-fold change. Readthrough was measured between 0 and 10 kb downstream from genes. n=2124 transposons in (A) and n=6061 transposons in (B) included in the analysis.

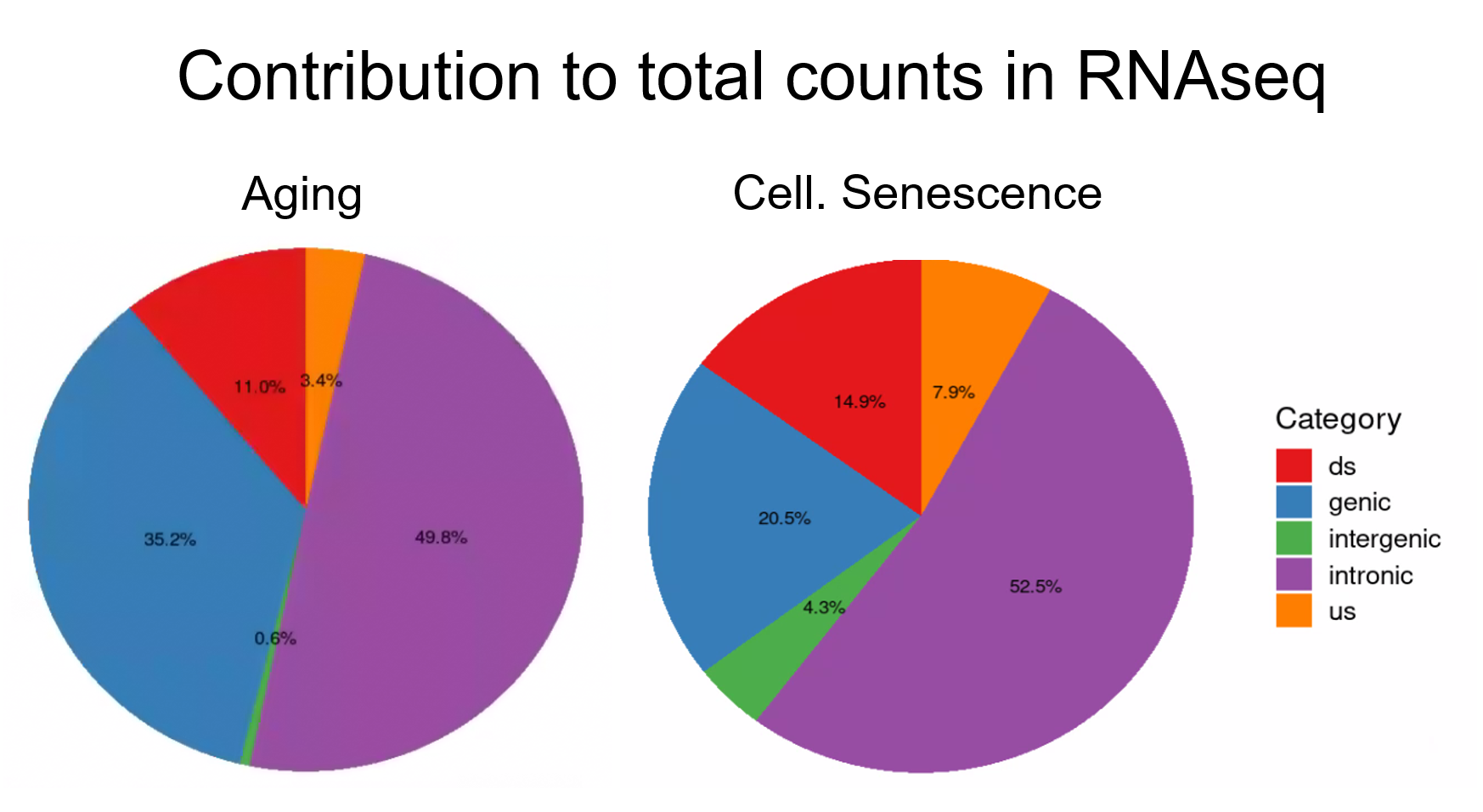

**Fig. S16. Intergenic transposons are rarely expressed.**
Total counts are the sum of all counts from transposons located in introns, genes, downstream (ds) or upstream (us) of genes (distance to gene < 25 kb) or in intergenic regions (distance to gene > 25 kb). Counts were defined as cumulative counts across all samples.

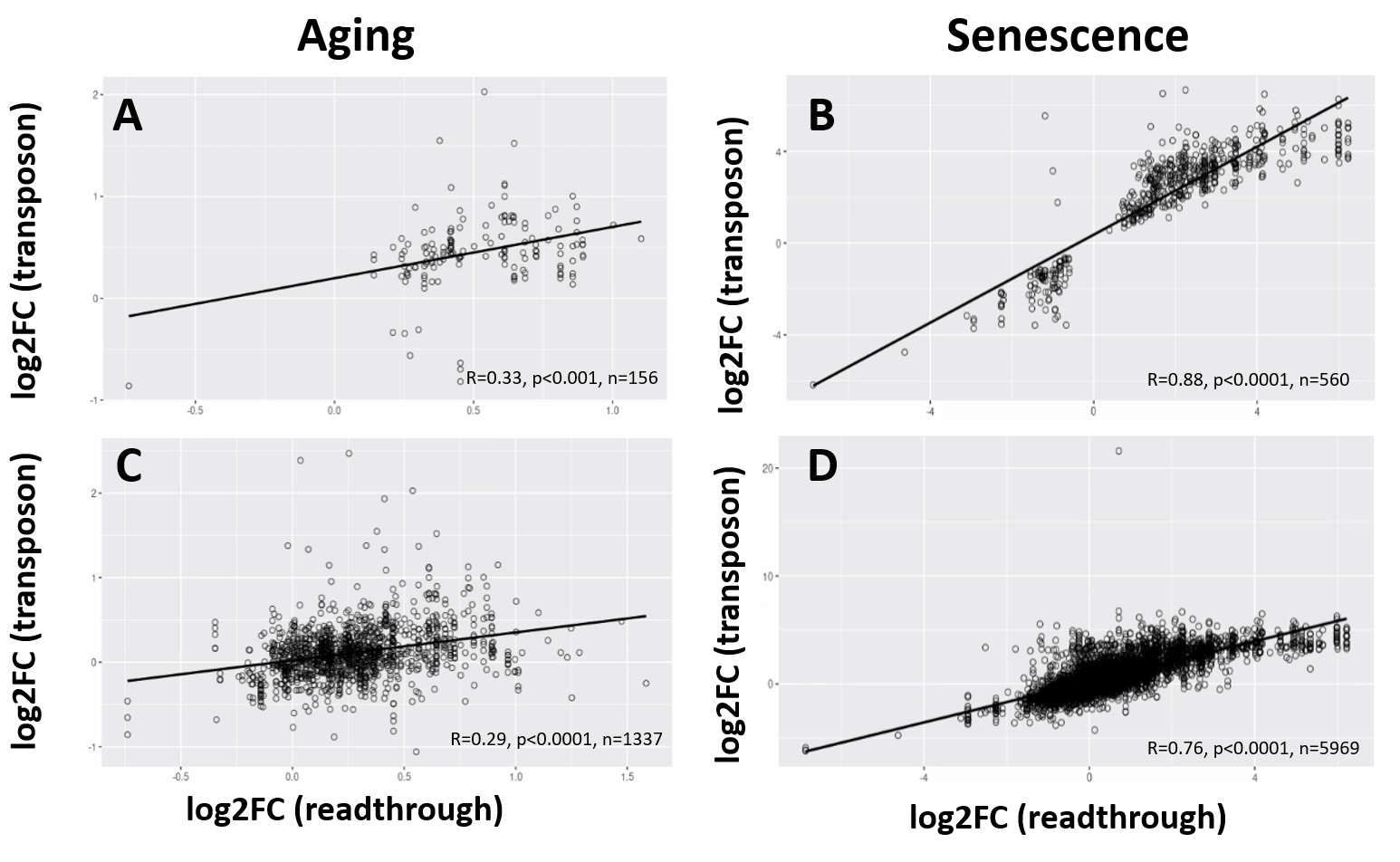

**Fig. S17. Age-related changes in readthrough are correlated with changes in transposon expression.**
Readthrough and transposon expression are correlated. For every transposon downstream of a gene, the correlation between the log-fold change (logFC) of the transposon and the adjacent readthrough region is plotted. Readthrough was measured in a 10 to 20kb region downstream of the gene.
A) Log-fold changes for all significant transposons plotted against log-fold changes for all significant, adjacent readthrough regions in the aging dataset.
B) Log-fold changes for all significant transposons plotted against log-fold changes for all significant, adjacent readthrough regions in the senescence dataset.
C) Log-fold changes for all transposons plotted against log-fold changes for all adjacent readthrough regions in the aging dataset.
D) Log-fold changes for all transposons plotted against log-fold changes for all adjacent readthrough regions in the senescence dataset.

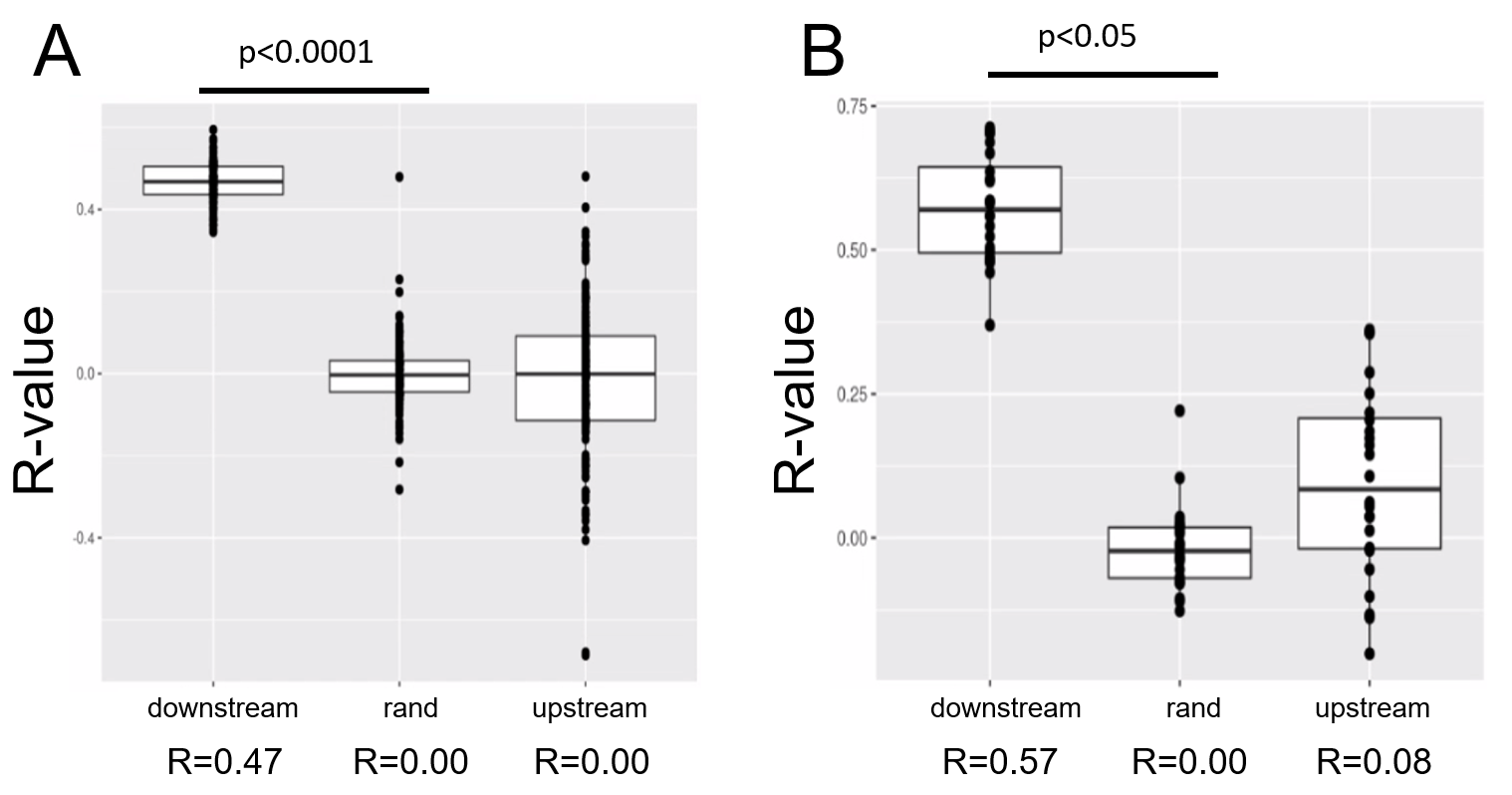

**Fig. S18. Positive correlation between read-through counts and transposons downstream of genes.**
Readthrough counts are correlated with the counts of transposons downstream of genes in the aging dataset (A) and in the senescence dataset (B). In contrast, readthrough counts do not correlate with counts of transposons upstream of genes. Readthrough counts also show no correlation with randomized transposon counts (labeled “rand”). Pearson correlation was performed for each sample in these datasets and the R-values for all the samples are shown here. Readthrough and transposons were restricted to a region 10kb upstream or downstream of genes.

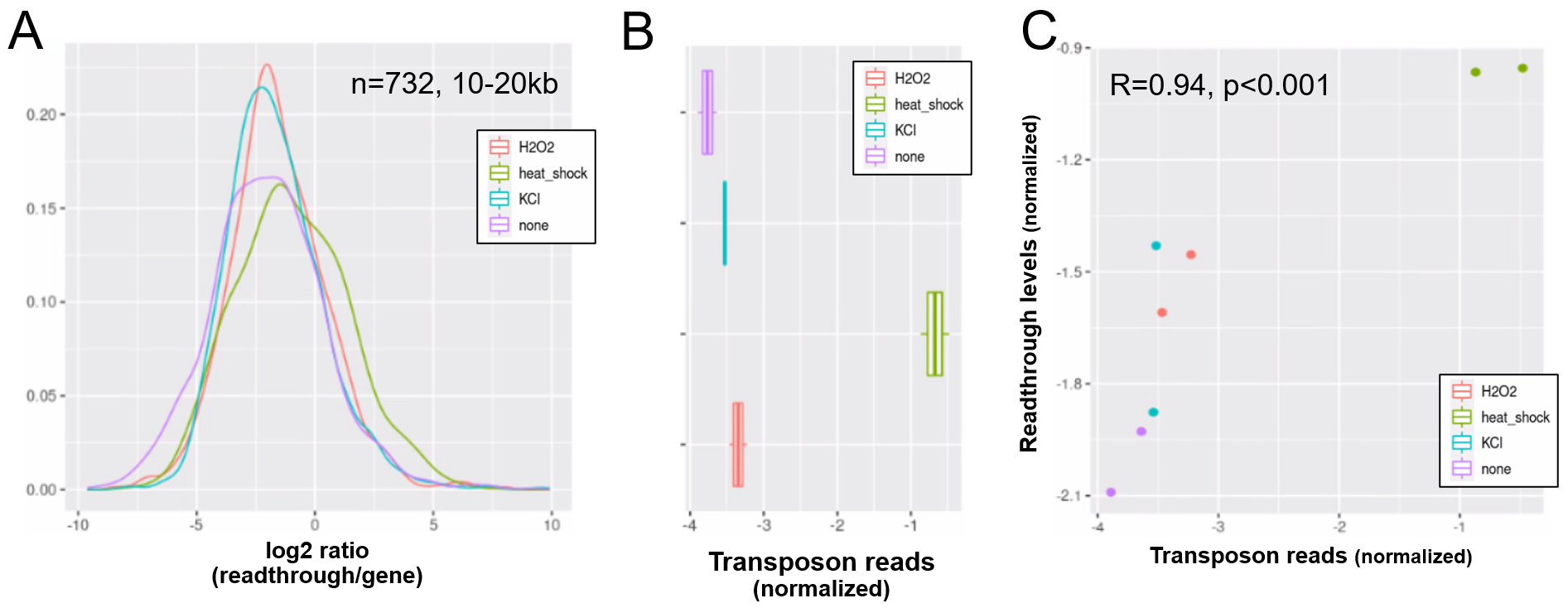

**Fig. S19. Elevated transposon expression after induced readthrough.**
Readthrough transcription is increased after heatshock treatment of NIH-3T3 fibroblasts and to a lesser extend after KCl and hydrogen peroxide (A). These readthrough inducing treatments also promote increased transposon expression (B). Therefore readthrough and transposon expression are correlated on a per-sample-basis (C). Data from **Vilborg et al. (2017).**
A) Readthrough was determined in a region 10 to 20kb downstream of genes for a subset of genes that was at least 20kb away from the nearest neighbouring gene (n=732 genes).
B) To normalize transposon expression counts for each transposon were corrected for the expression of the nearest gene. Normalized transposon counts for each sample are shown as box-whisker plot.
C) Normalized transposon counts (as in B) and readthrough counts (medians) are plotted for each sample (n=8).

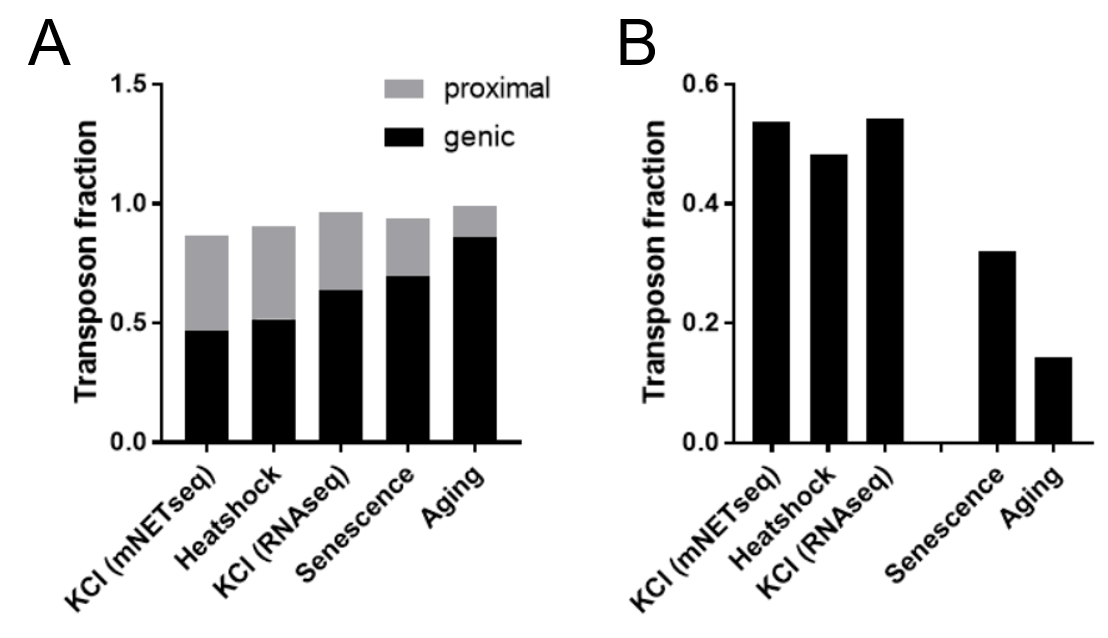

**Fig. S20. Heatshock and osmotic stress promote expression of gene proximal transposons.**
Specific readthrough induction leads to stronger differential expression of gene proximal transposons (A) and stronger upregulation of extragenic transposons (B) as compared with cellular senescence and aging, which are associated with more modest readthrough.

Differential expression was performed for KCl treated HEK293 cells vs control (mNETseq; Bauer et al. 2018), 44°C heatshocked NIH-3T3 vs control (RNAseq; Vilborg et al. 2017), KCl treated HEK293 cells vs control (RNAseq; Rosa-Mercado et al. 2021), senescent cells and aged fibroblasts (as detailed in Fig. 1). The fraction of all genic and gene proximal transposons (distance < 25kb) among differentially regulated transposons is shown in (A); the remaining transposons are intergenic (distance > 25kb). The fraction of all extra-genic transposons among the significantly upregulated transposons is shown in (B).
